# Multidimensional Data Organization and Random Access in Large-Scale DNA Storage Systems

**DOI:** 10.1101/743369

**Authors:** Xin Song, Shalin Shah, John Reif

## Abstract

With impressive density and coding capacity, DNA offers a promising solution for building long-lasting data archival storage systems. In recent implementations, data retrieval such as *random access* typically relies on a large library of non-interacting PCR primers. While several algorithms automate the primer design process, the capacity and scalability of DNA-based storage systems are still fundamentally limited by the availability of experimentally validated orthogonal primers. In this work, we combine the nested and semi-nested PCR techniques to virtually enforce multidimensional data organization in large DNA storage systems. The strategy effectively pushes the limit of DNA storage capacity and reduces the number of primers needed for efficient random access from very large address space. Specifically, our design requires *k* * *n* unique primers to index *n*^*k*^ data entries, where *k* specifies the number of dimensions and *n* indicates the number of data entries stored in each dimension. We strategically leverage forward/reverse primer pairs from the same or different address layers to virtually specify and maintain data retrievals in the form of rows, columns, tables, and blocks with respect to the original storage pool. This architecture enables various random-access patterns that could be tailored to preserve the underlying data structures and relations (*e.g*., files and folders) within the storage content. With just one or two rounds of PCR, specific data subsets or individual datum from the large multidimensional storage can be selectively enriched for simple extraction by gel electrophoresis or readout *via* sequencing.

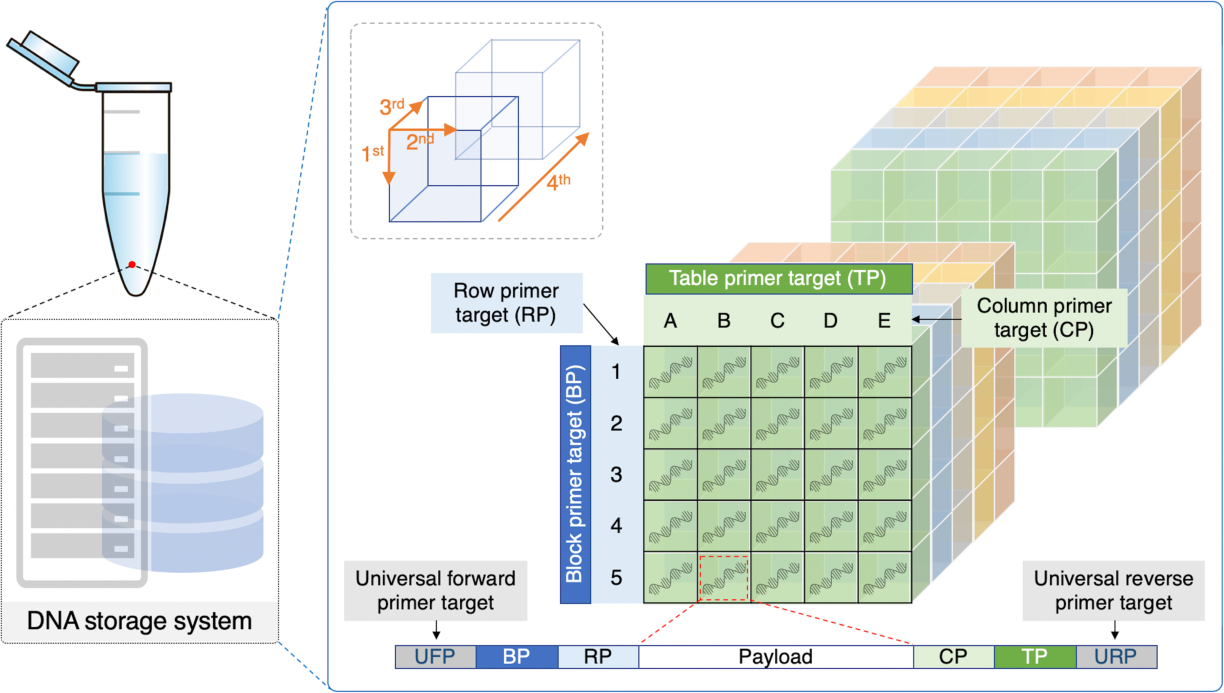

---

Advances in DNA synthesis and sequencing technologies have allowed researchers to explore digital data storage in synthetic DNA molecules.^1^ Several DNA archival storage systems have been recently constructed and supported PCR-based random access for data retrieval.^2–5^ The data is typically segmented into chunks to encode as oligonucleotides (oligos) with fixed lengths appropriate for the current DNA synthesis and sequencing platforms. As a result, data retrieval and reconstruction must rely on address sequences appended to individual data-encoding oligos in the storage pool such that PCR reactions could utilize these addresses as primer targets to selectively amplify the desired strands from a large pool of oligos. To ensure the specificity of random access in large DNA databases, a vast number of orthogonal primers need to be meticulously designed to reduce cross-talking and mispriming. This type of addressing mechanism becomes unscalable and unreliable as the storage content increases in size, posing a major bottleneck for implementing practical DNA-based storage systems at large economical scales. A recent exciting work^2^ demonstrated individual file retrieval from a DNA storage pool encoding over 200 MB of data. Oligos belonging to the same file shared the same PCR primer targets and relied on an additional unique address to reassemble the sequences into the correct order within a file. With error-correcting codes, the system supported reliable random access at the single file level. However, this implementation could not be used to explore and provide different data retrieval patterns. Specifically, retrieving groups of related files would require the same number of unique primers as compared to retrieving those files one by one, which is costly and inefficient. In this letter, we investigate the designs of multidimensional DNA storage systems with a scalable architecture that combines nested and semi-nested PCR techniques, both of which are simple methods supporting highly sensitive and specific amplification of target oligos from very large and noisy background.^6^ We illustrate that such architecture could not only efficiently scale up PCR-based random access by using a relatively small number of primers but also give rise to multiple well-defined data retrieval patterns, which may be strategically leveraged to organize and access large DNA databases in simple formats such as rows, columns, tables, and blocks. In theory, our proposed design could leverage the efficient use and reuse of *k* * *n* orthogonal primers to establish a virtual address system for DNA storage of size *n*^*k*^. As an example, we illustrate arbitrary indexing within an *n*^4^ space using only 4 * *n* orthogonal primers, and every random-access operation (including the retrieval of single data entries) demonstrated here can be accomplished with no more than two rounds of PCR. These operations maybe be extended with multiplex PCR to concurrently enrich several arbitrarily selected data subsets from the storage. Our design adheres to standard experimental protocol for nested and semi-nested PCR^6^ and therefore supports simple data extractions *via* sequencing or gel electrophoresis without sophisticated physical separation of oligos.

Nested PCR is a well-established technique for improving the specificity and sensitivity of PCR reactions, where the amplicon from the first PCR reaction serves as the template for the second PCR amplification primed by an inner primer pair.^6^ Nested PCR was previously studied in the context of building hierarchical DNA memories. Kashiwamura *et al*.^7^ appended three address blocks on one end of the data-encoding oligos and used a common reverse primer target on the opposite end. Access to a single data oligo was attained by specifying the sequential order of the address primers. The design concept was analogous to semi-nested PCR,^8^ where one of the outer primers used for the first round of PCR is also used as an inner primer during the second PCR amplification. Semi-nested PCR was also employed in a recent work by Tomek *et al*.^9^ to improve the DNA storage capacity by combining a simple two-primer nested address system with physical separations for target file enrichment. Yamamoto *et al*.^10^ expanded the nested primer hierarchy and concatenated multiple primer targets on both ends of the data oligos. Each primer pair was referred to as an address layer, and data retrieval operations started with priming the outer-most layer towards the inner layers. Their work experimentally demonstrated that by specifying a unique primer pair for each address layer, the target oligo could be amplified and extracted from an enormous address space after several rounds of nested PCR. However, their primary focus was to scale up the nested primer hierarchy and demonstrate the specificity of single target oligo retrieval. In contrast, our design goal is threefold: (i) providing a systematic architecture to enforce virtual organization of data oligos in a scalable and multidimensional address space, (ii) minimizing the number of orthogonal primers needed for uniquely indexing entries from a large storage pool, and (iii) strategically combining nested and semi-nested PCR (*i.e*., allowing the use of forward/reverse primers from the same or different address layers during data retrievals) to support several well-defined random-access patterns in large DNA databases. Compared to prior works, the amplicon after each PCR round (including the intermediate rounds) could be consistently viewed as meaningful random-access result from our DNA storage, because the primer pairs are always specified to enforce and maintain useful relations such as rows, columns, tables, or blocks in the resulting PCR product with respect to the original storage pool. As a result, DNA database designers may be able to take advantage of such architecture and tailor the different random-access patterns to the underlying data relations that may already exist in their storage content. Moreover, our design strategy could be easily modified to allow different configurations of data organization for multidimensional DNA storage. Here we illustrate two design variations, where the first design supports 16 random-access patterns at the block/table/row/column/entry levels *via* one or two rounds of PCR, and the second design eliminates the block-level data organization to simplify several particularly useful random-access patterns *via* only single-round PCR. We do not discuss the details on experimental implementations of DNA storage systems, which involve other important aspects such as data encoding, error correction, and system integration and automation.^11–14^

In design A (Figure 1), the pool of data oligos is virtually organized in the form blocks, tables, rows, and columns. Each data-encoding strand is composed of several domains, including a data payload block surrounded by three address blocks (*i.e*., PCR primer targets) on both sides. These address blocks are named and arranged based on their specific roles in establishing the different data organization levels and random-access patterns. For example, a four-dimensional DNA storage can contain multiple 3D data blocks, and each block can contain multiple 2D data tables. Blocks are distinguished by different BP (block primer target) and the same set of TPs (table primer target) are reused across all blocks. Tables in the same block share the same BP and differ in TP. All entries in a given table share the same BP/TP pair. RP (row primer target) identifies the row, and CP (column primer target) identifies the column. The same set of RP/CP pairs are reused across all tables. All oligos share the same UFP (universal forward primer target) and URP (universal reverse primer target) pair to support data retrieval from all blocks. Pairing an inner address block with UFP or URP enables additional useful data retrieval patterns. The number of orthogonal primers needed to index arbitrary entries in this four-dimensional address space is calculated in Table 1. Approximately, this architecture uses 4 * *n* orthogonal primers to uniquely index *n*^4^ data entries. Figure 2 illustrates 16 different patterns of data retrieval enabled by this architecture. Depending on the retrieval pattern, the target data subset is enriched *via* either one or two rounds of PCR. It is worth noting that when the target oligos are simultaneously enriched from multiple addresses, several random-access patterns could maintain the mutual distinguishability of retrieved data oligos across all address layers (*i.e*., the addresses of enriched oligos can be inferred from their remaining inner-layer primer targets), and therefore further data reassembly can be carried out *in silico* once the extracted oligos are sequenced. For patterns that lose the mutual distinguishability among the retrieved oligos, the respective data retrieval operations could still be useful for applications that cares about the storage content but not the ordering of the retrieved data oligos.

**Table 1.** Total number of orthogonal primers and random-accessible entries in design A.

|  |
| --- |
| *Number of orthogonal PCR primers needed ($4 * n$)* |
| $= (\# \text{ of rows in a table}) + (\# \text{ of columns in a table}) + (\# \text{ of tables in a block}) + (\# \text{ of blocks}) + 2$ |
| *Number of unique data entries addressable ($n^4$)* |
| $= (\# \text{ of rows in a table}) \times (\# \text{ of columns in a table}) \times (\# \text{ of tables in a block}) \times (\# \text{ of blocks})$ |

**Figure 1.**
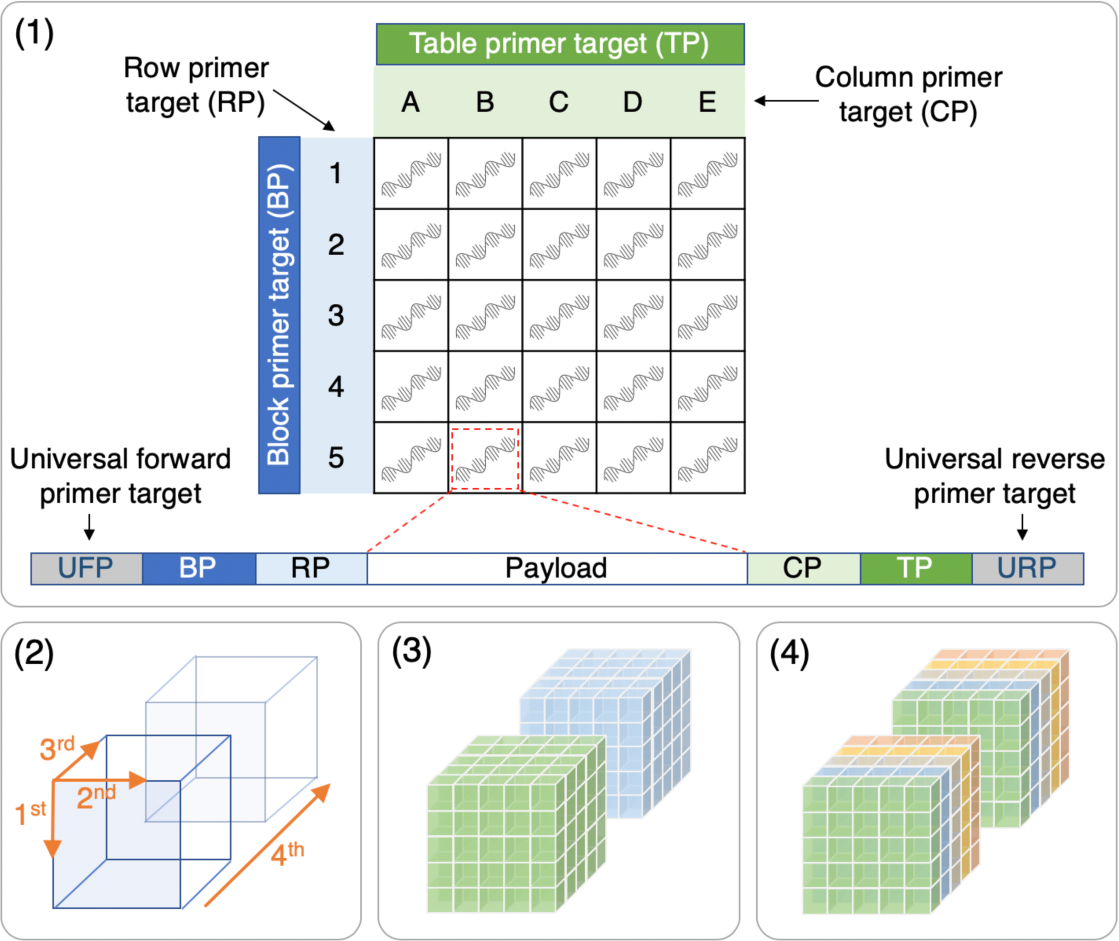
Multidimensional data organization in design A. (1) Design of data-encoding strands and their organization as addressable entries in a 2D table. (2) Illustration of a four-dimensional addressable space. The 1^st^ to 4^th^ dimensions refer to rows, columns, tables, and blocks, respectively. (3) Data blocks are distinguishable by primer target BP. (4) Data tables within each block are distinguishable by primer target TP.

**Figure 2.**
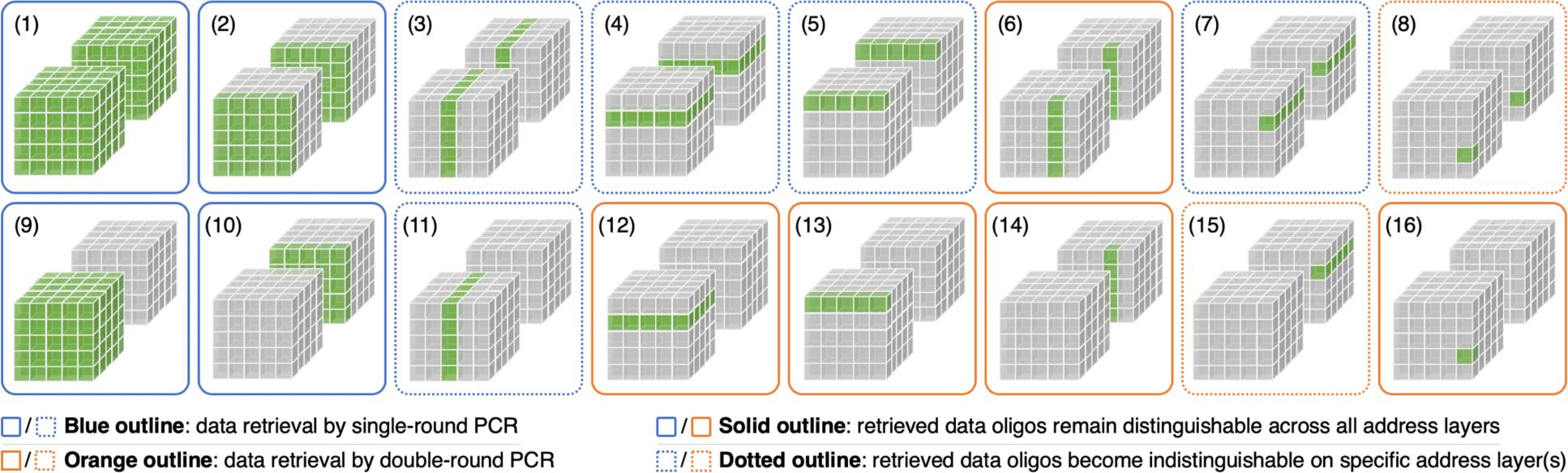
Data retrieval patterns supported by design A. **(1)** UFP+URP: all blocks. **(2)** UFP+TP: specific table from all blocks. **(3)** UFP+CP: specific column from all tables in all blocks. **(4)** RP+URP: specific row from all tables in all blocks. **(5)** RP+TP: specific row from specific table in all blocks. **(6)** UFP+TP then UFP+CP: specific column from specific table in all blocks. **(7)** RP+CP: specific entry from all tables in all blocks. **(8)** UFP+TP then RP+CP: specific entry from specific table in all blocks. **(9)** BP+URP: specific block. **(10)** BP+TP: specific table from specific block. **(11)** BP+CP: specific column from all tables in specific block. **(12)** BP+URP then RP+URP: specific row from all tables in specific block. **(13)** BP+TP then RP+TP: specific row from specific table in specific block. **(14)** BP+TP then BP+CP: specific column from specific table in specific block. **(15)** BP+URP then RP+CP: specific entry from all tables in specific block. **(16)** BP+TP then RP+CP: specific entry from specific table in specific block.

Here we show that the design A can be slightly modified for applications that do not need the block-level organization but require rapid random access to an arbitrary row/column from an arbitrary table *via* single-round PCR. In design B (Figure 3), each entry in the table corresponds to a data-encoding strand composed of several domains. Tables can be distinguished by either TP1 or TP2. All entries in a given table share the same TP1/TP2 pair. RP identifies the row, and CP identifies the column. The same set of RP/CP pairs are reused across all tables. The same UFP/URP pair is used as the outer address layer on all data oligos to enable simultaneous retrieval of all tables or a specific row/column from all tables through the RP/URP or UFP/CP pair. The number of orthogonal primers needed to index arbitrary entries in this three-dimensional address space specified by design B is calculated in Table 2. Approximately, this architecture uses 3 * *n* orthogonal primers to uniquely index *n*^4^ data entries. Figure 4 illustrates 8 different data retrieval patterns enabled by this architecture along with 11 extended retrieval patterns (2a to 8a, 5b to 8b) by incorporating multiplex PCR. Except for single-entry random access, all the other data retrieval patterns operate on single-round PCR.

**Table 2.**
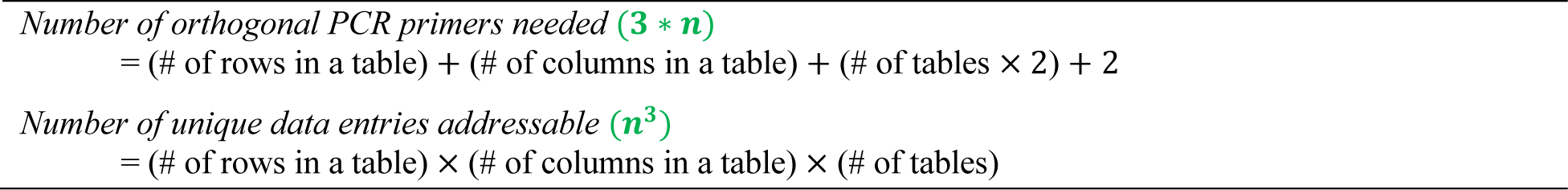
Total number of orthogonal primers and random-accessible entries in design B.

**Figure 3.**
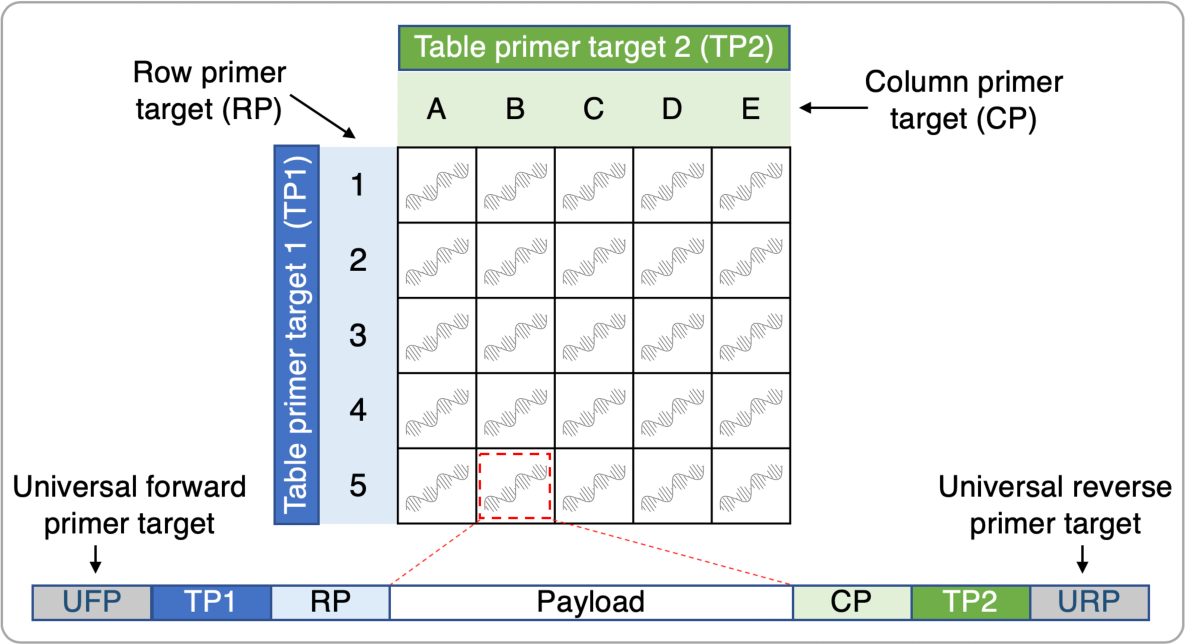
Multidimensional data organization and design of data-encoding strands in design B.

**Figure 4.**
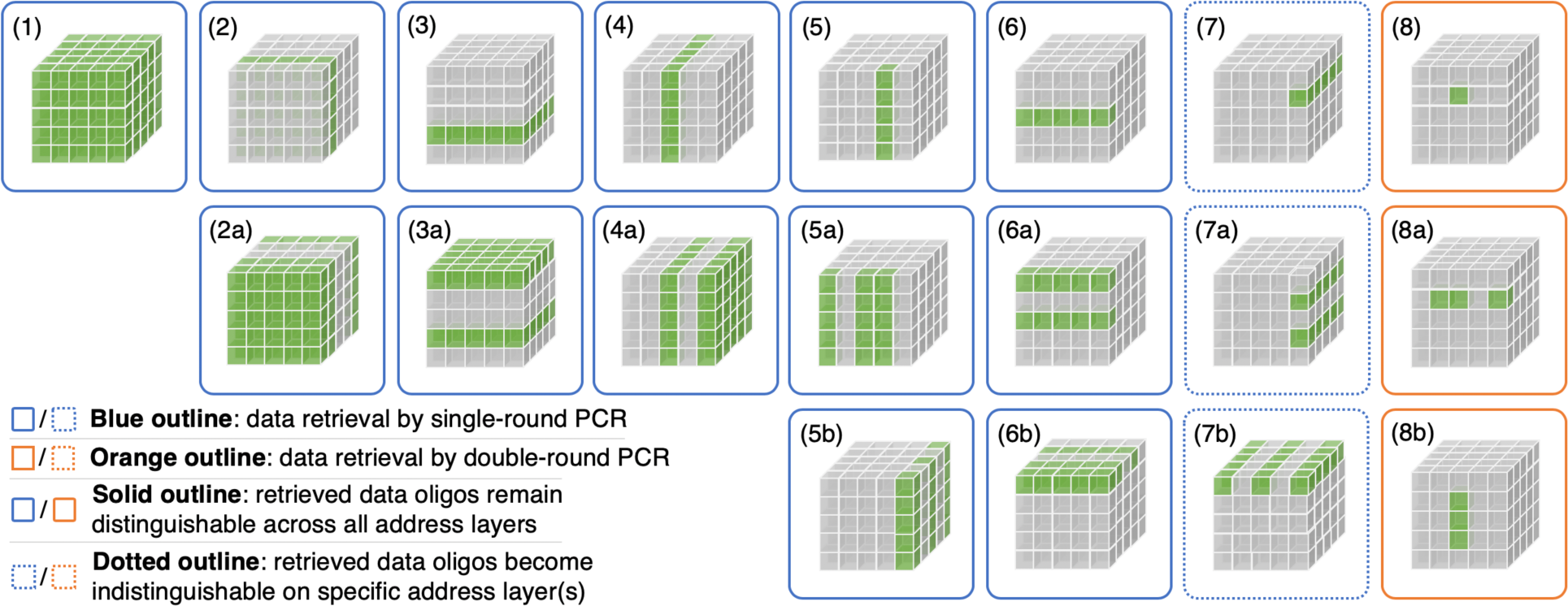
Data retrieval patterns supported by design B. **(1)** UFP+URP: entire database. **(2)** TP1+TP2 (or TP1+URP or UFP+TP2): specific table. **(2a)** TP1(multiplex)+URP: multiple tables. **(3)** RP+URP: specific row from all tables. **(3a)** RP(multiplex)+URP: multiple rows from all tables. **(4)** UFP+CP: specific column from all tables. **(4a)** UFP+CP(multiplex): multiple columns from all tables. **(5)** TP1+CP: specific column from specific table. **(5a)** TP1+CP(multiplex): multiple columns from specific table. **(5b)** TP1(multiplex)+CP: specific column from multiple tables. **(6)** RP+TP2: specific row from specific table. **(6a)** RP(multiplex)+TP2: multiple rows from specific table. **(6b)** RP+TP2(multiplex): specific row from multiple tables. **(7)** RP+CP: specific entry from all tables. **(7a)** RP(multiplex)+CP: entry on specific column from multiple rows from all tables. **(7b)** RP+CP(multiplex): entry on specific row from multiple columns from all tables. **(8)** TP1+TP2 then RP+CP: specific entry from specific table. **(8a)** TP1+TP2 then RP+CP(multiplex): multiple entries on specific row from specific table. **(8b)** TP1+TP2 then RP(multiple)+CP: multiple entries on specific column from specific table.

In this work, we have introduced a strategy that combines nested and semi-nested PCR to organize large multidimensional DNA data storage systems while significantly reducing the number of orthogonal primers needed for supporting efficient random access in multiple patterns. A recent algorithm^2^ was proposed to design up to 14000 pairs of orthogonal 20-mer primers for PCR-based random access in large DNA storage pools, which could potentially scale up to archive a few terabytes (TBs) of digital data. With our design strategy, we expect that the same number of primers could be efficiently utilized to create an enormous address space and dramatically increase the storage capacity. Here we illustrate with an example based on our design in Figure 1. Suppose we take 2 primers from the 28000-primer library for UFP and URP and then evenly distribute the remaining primers for use with BP, TP, RP, and CP. This would allow the construction of a very large DNA storage containing roughly 7000 blocks, each of which consists of 7000 tables with 7000 rows and 7000 columns in each table (*n* = 7000, *k* = 4). As a result, the architecture would allow 7000^4^ (∼ 2401 trillion) data oligos to be stored, organized, and uniquely indexed. If we assume that the length of each oligo is 200 nucleotide (nt) and the address blocks are 20-mers, then every oligo in our design could carry a data payload of 80 nt (200 − 20 * 6). While in theory each nucleobase may encode up to 2 bits, we assume a coding density of 1 bit/nt to account for the coding redundancies needed for error correction. Accordingly, each oligo would encode 80 bits, which amounts to 24010 TB (80 * 7000^4^ bits) of digital data in the whole storage pool. Remarkably, each 80-bit data chunk in this 24-petabyte storage is uniquely addressable and retrievable by two simple PCR rounds *via* the random-access pattern (16) illustrated in Figure 2. Theoretically, the storage capacity would expand further as the DNA synthesis and sequencing technologies improve such that the data could be encoded on longer synthetic oligos at a higher coding density. Note that the 1 bit/nt assumption we made above was rather conservative since recent works have already experimentally demonstrated higher coding densities in DNA storage, as summarized in our latest review.^1^ Moreover, our proposed design facilitates virtual organization of the data-encoding oligo pool as a scalable and multidimensional storage architecture that readily supports different useful patterns of data retrievals. For example, while each 80-bit data segment is viewed as an “entry” in the storage space, a group of data segments that form a “file” could be collectively stored as a table. As multiple tables are organized in a block, the block level could be effectively used for organizing data in “folders”. To efficiently utilize the enormous address space supported by our architecture, the size of tables and blocks could be flexibly tailored to the data content, and techniques such as tail merging may be adapted from computer file system design to mitigate data allocation issues such as fragmentation in the DNA storage. While we only demonstrated designs for three- and four-dimensional data organization, our proposed architecture could be readily scaled up with additional dimensions by adding primer targets on the oligos. Although the strategic combination of nested and semi-nested PCR allows very efficient reuse of primers on the inner address layers (*i.e*., oligos organized in different 5^th^-dimension groups could share the same set of primers for the 4^th^-dimension indexing, and so on), concatenating additional primer targets on oligos would shorten the effective payload length and diminish the overall coding density of the DNA storage pool. Furthermore, single-entry random access would likely require more than two rounds of PCR if the storage is extended beyond 4 dimensions. On smaller scales, our design may also find useful implementations such as DNA-based lookup tables (*e.g*., encoding numerical weights or Boolean values of useful functions) to potentially interface with DNA computing for interesting applications. Other complex systems of DNA databases may be enabled by encoding entries in one database as “pointers” that establish particular relational networks with data oligos from another DNA database and so forth.

## Acknowledgments

This work was sponsored by NSF grant no. CCF 1617791. X.S. acknowledges support from NSF grant no. DGE 1545220. We would like to thank Dr. Joshua Granek and Xiangyu Zhang for helpful discussions.

